# Prolonged thermal stress enhances mosquito tolerance to viral infection

**DOI:** 10.1101/2024.09.06.611661

**Authors:** Hugo D. Perdomo, Ayda Khorramnejad, Nfamara M. Cham, Alida Kropf, Davide Sogliani, Mariangela Bonizzoni

## Abstract

How and to what extent mosquito-virus interaction is influenced by climate change is a complex question of ecological and epidemiological relevance. We worked at the intersection between thermal biology and vector immunology and studied shifts in tolerance and resistance to the cell fusing agent virus (CFAV), a prominent component of the mosquito virome know to contribute to shaping mosquito vector competence, in warm-acclimated and warm-evolved *Aedes albopictus* mosquitoes. We show that the length of the thermal challenge influences the outcome of the infection with warm-evolved mosquitoes being more tolerant to CFAV infection, while warm-acclimated mosquitoes being more resistant and suffering from extensive fitness costs. These results highlight the importance of considering fluctuations in vector immunity in relation to the length of a thermal challenge to understand natural variation in vector response to viruses and frame realistic transmission models.

## Introduction

Current anthropogenic climate change has profound and complex implications for the prevalence and the transmission dynamics of arboviruses such as dengue, Zika and chikungunya, which are an impending risk for 3.9 billion people globally^1,2^. Climate warming is expected to cause large shifts in the distributions of the main arboviral vectors, the mosquitoes *Aedes aegypti* and *Aedes albopictus*, and changes in their phenology, because mosquitoes are ectotherms, and environmental temperature (T_*a*_) impacts nearly all aspects of their life cycle ^3,4^. Increase in T_*a*_ is also known to accelerate viral replication in mosquitoes ^5^, but whether mosquito response to viruses is counter-modulated by T_*a*_ remains largely unexplored.

Organism can respond to a viral infection by actively trying to limit viral replication, a strategy called resistance, or by controlling the cost of infection without impeding viral replication, a strategy called tolerance ^6^. Tolerance is not well-understood in mosquitoes because to date vector immunology has been biased by an anthropogenic perspective and focused on identifying pathways used by mosquitoes to resist viral infections to leverage them into transmission control strategies ^7,8^. Resistance and tolerance can act independently, synergistically or alternating, but have different ecological and evolutionary repercussions ^9-12^.

By directly limiting viral replication, resistance mechanisms impose a strong selective pressure on viruses ^13,14^. In contrast, tolerance minimises cost of infection, resulting in neutral or positive effects on pathogens ^15^. Understanding whether global climate change favours a shift between resistance and tolerance, or their synergistic interaction, is crucial because the efficiency of one immunological response over the other has immediate effects on viral transmission dynamics and long-term impacts on the evolution of both vectors and viruses ^16-18^.

In this study we used *Ae. albopictus* and the orthoflavivirus cell fusing agent virus (CFAV, *Flaviviridae* family) to explore if T_*a*_ alters the dynamic response of mosquitoes to a viral infection. *Aedes albopictus* is an invasive species originally from southeast Asia, which established in both tropical and temperate areas of the word and emerged as the main arboviral vector in Europe and North America ^19^. CFAV is the first-identified and, so far, the best-characterised *Aedes* insect-specific virus (ISV) ^20-26^. ISVs are insect-restricted viruses, which are found with higher prevalence and frequency than arboviruses in field mosquitoes ^27-29^. Additionally, ISV infection is known to elicit mosquito immunity ^30^, thus alerting vector competence and contributing to the natural heterogeneity of arboviral transmission dynamics ^31^. Despite having direct implications for mosquito biology and indirect effects on the efficiency and flux of arboviral transmission, the impact of T_*a*_ on the relationship between ISVs and mosquitoes remains largely unexplored.

We also reasoned that anthropogenic climate change manifests not only through global warming, but also through extensive T_a_ fluctuations, which can last few days or months ^32,33^. To test if immunological mechanisms induced by a short-term thermal challenge differ from those resulting from a constant and prolonged exposure to heat ^32,33^, we studied CFAV infection in mosquitoes exposed to 32°C, a temperature above the optimum for *Ae. albopictus* ^3^, for one or ten generations.

We show that exposure to heat increases overall *Ae. albopictus* tolerance to CFAV infection, but the length of the thermal challenge alters mosquito fitness and influence the immunological response to CFAV. Mosquitoes exposed to heat for one generation fight CFAV infection, whereas mosquitoes conditioned to heat through several generations are healthier than those not exposed to a thermal challenge and tolerate CFAV infection.

## Results

We exposed *Ae. albopictus* mosquitoes to a hot thermal regime (32°C for 14 h and 26°C for 10 h) for one or ten generations generating warm-acclimated and warm-evolved mosquitoes, respectively (Fig. 1). We infected these mosquitoes with CFAV and studied immunological traits of tolerance and resistance, as well as the cost of infection in terms of mosquito longevity and its reproductive output in comparison to infection of mosquitoes maintained under our standard laboratory thermal conditions, hereon referred to as “standard”.

**Figure 1.**
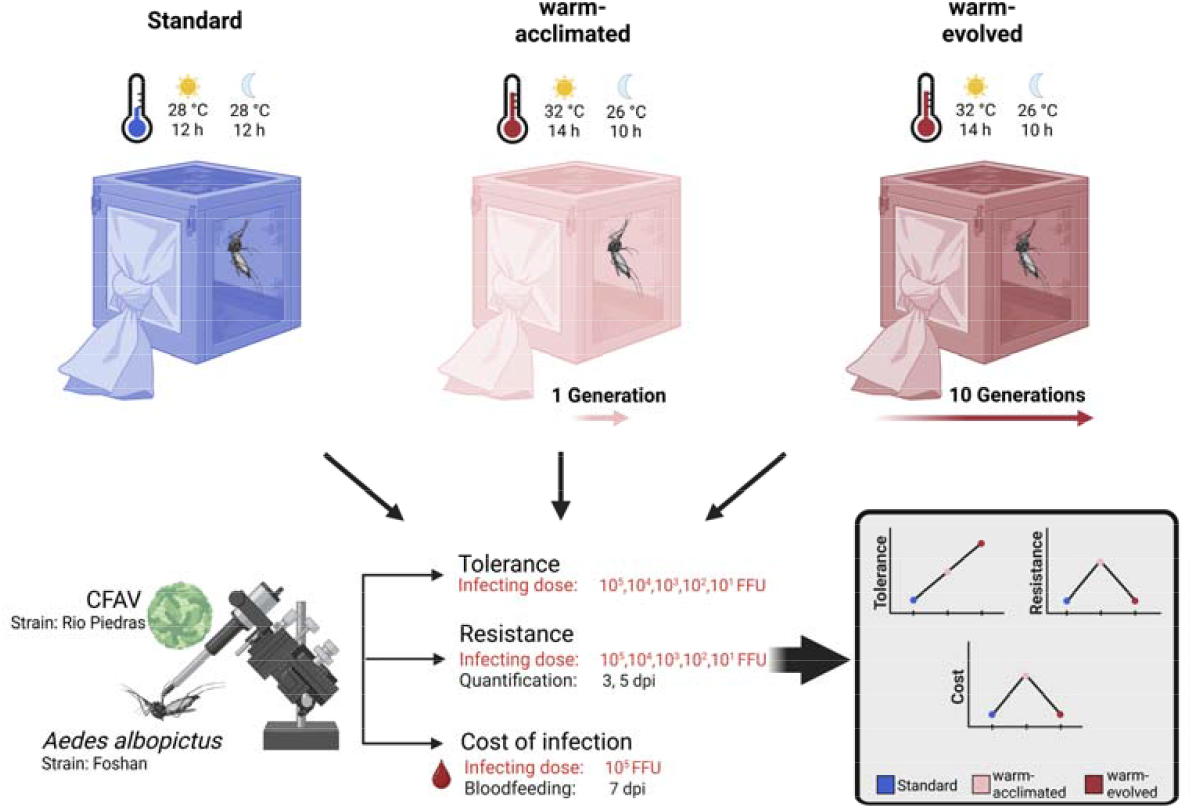
Schematic representation of the experimental design and results. We compared tolerance and the resistance to CFAV and the fitness of CFAV infected mosquitoes maintained under a hot (32°C for 14 h and 26°C for 10 h) thermal regime or our standard laboratory conditions (constant 28°C). Mosquitoes were exposed to the hot thermal regime for either one or ten generations resulting in warm-acclimated and warm-evolved mosquitoes, respectively. Representative results of resistance, tolerance and mosquito fitness are shown in the bold frame.

### Exposure to hot increases viral tolerance of *Ae. albopictus* mosquitoes

We measured tolerance as the rate of change in host fitness at increasing viral loads as recently proposed ^11,34^. To do so, we infected *Ae. albopictus* mosquitoes with five doses of CFAV spanning 10 to 10^5^ viral particles and followed mosquito survival after infection. We saw reduced longevity in mosquitoes maintained at standard conditions when we compared mock infected vs CFAV-infected mosquitoes, starting with 10^2^ viral particles (Fig. 2A, Table S1). We observed that the longevity of warm-acclimated mosquitoes was significantly reduced with respect to mock-infected mosquitoes only after infection with the highest CFAV dose (Media−10^5^ viral particles, z.ratio=-3.493, Df=inf, p=0.0063). We did not see significant changes in longevity of warm-evolved mosquitoes at any CFAV infecting doses (Fig. 1 B, C, Table S1). We found that the regimen at which mosquitoes were reared (Chisq=8.762, Df=2, p=0.013), the injected viral dose (Chisq=63.381, Df=5, p=2.43 x 10^−12^), and their interaction reduced survival (Chisq=21.851, Df=10, p=0.016).

**Figure 2.**
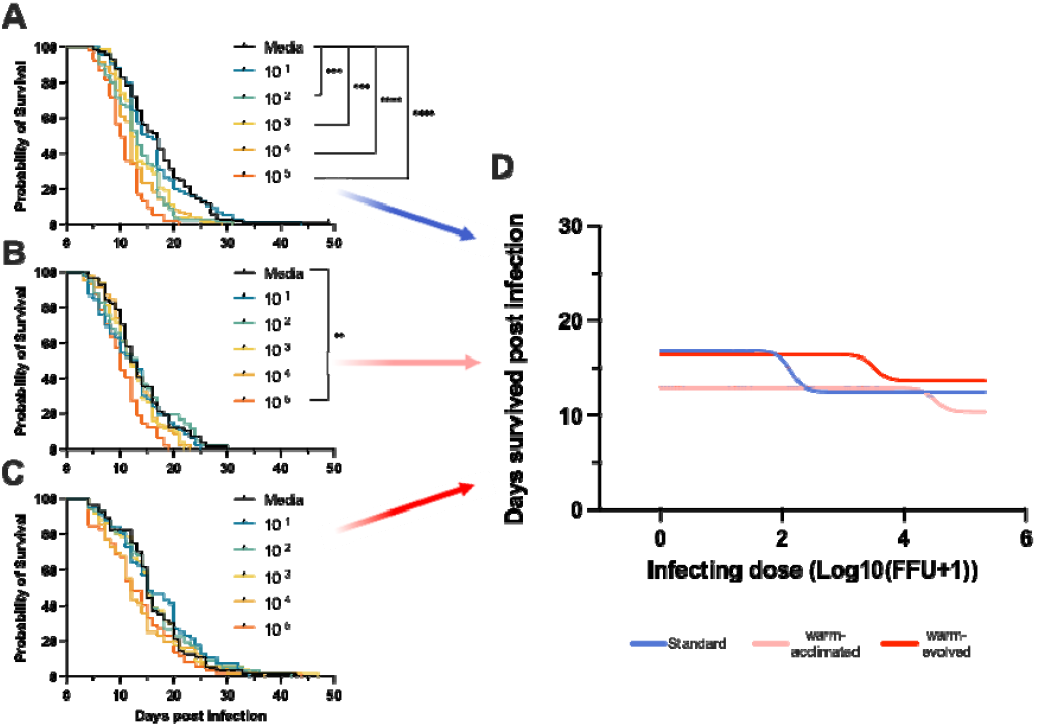
Exposure to hot increases *Ae. albopictus* tolerance to CFAV infection. Survival probability of (A) mosquitoes kept at standard conditions, (B) warm-acclimated and (C) warm-evolved after mock infection (media) and infection with five different concentrations of CFAV (10^5^, 10^4^, 10^3^, 10^2^, 10 viral particles). In all graphs, we used the Kaplan–Meier survival analysis with a cox proportional hazard test to determine the effect of CFAV infection on mosquito survival. (D) Tolerance curves of warm-acclimated (pink), warm-evolved (red) and mosquitoes kept at standard conditions (blue) were built by plotting mosquito survival time after infection versus the log10 scale of infecting dose. Each point represents an individual mosquito, and lines are logistic fits of the data. Significant result as follows: ** p<0.01, *** p<0.001, **** p<0.0001.

**Figure 3.**
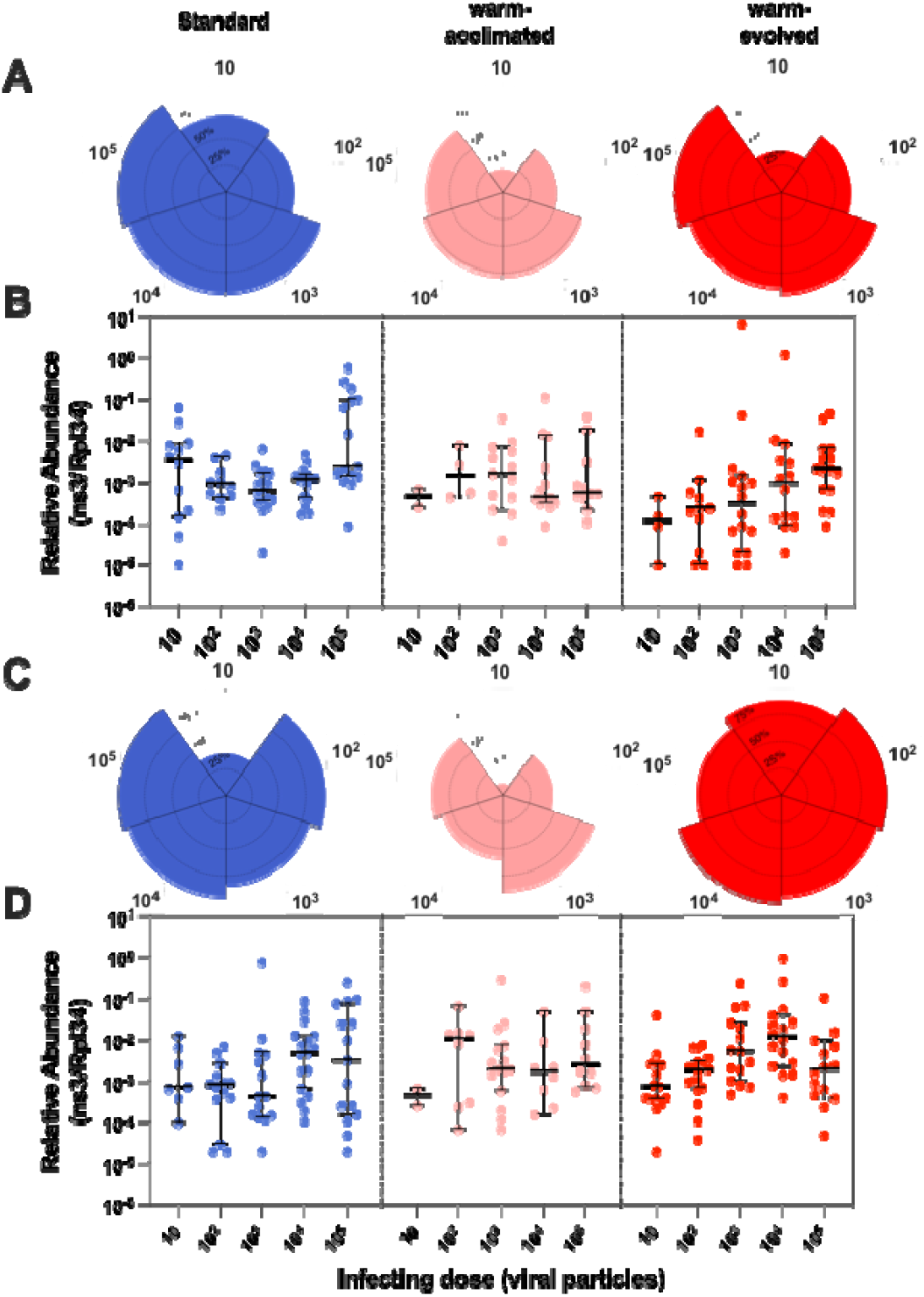
Acclimation to a hot temperature increases *Ae. albopictus* resistance. **(A)** Circular bar plot showing prevalence of infection as percentage of infected mosquitoes in relation to CFAV doses 3 dpi. (**B**) Quantification of CFAV genomes in relation to CFAV doses 3 dpi. (**C**) Circular plot showing prevalence of infection as percentage of infected mosquitoes in relation to CFAV doses 5 dpi. (**D**) Quantification of CFAV genomes in relation to CFAV doses 5 dpi. Data for warm-acclimated mosquitoes are in pink, warm-evolved mosquitoes in red and mosquitoes kept at standard condition in blue. In plots B and D, each point represents data from an individual mosquito; black solid lines and bars are the median of viral load and the 95% confidence intervals, respectively.

Next, we plotted the survival time of each mosquito in relation to the infecting CFAV dose and fitted a four-parameter logistic model with a Poisson regression to identify vigour, meaning the survival time of uninfected hosts; severity, meaning longevity of the host when infected with the highest viral load and sensitivity, or effective concentration 50 (EC_50_), which defines the viral load that reduces vigour by half ^35^. The choice of the model was based on the goodness of fit (Table S1) and previous studies ^35,36^. To make comparisons across thermal conditions, we also tried to constrain the slope of the curves to -1, as described elsewhere ^36^. The slope of the curve denotes the rate of loss of host health ^35^. We could not fit the tolerance curve of warm-acclimated mosquitoes with a slope to -1 (Table S1). This result indicates that, when summed to the challenge of infection, acclimation reduces mosquito health faster than what observed in both warm-evolved and mosquitoes maintained at standard condition. We found that observed differences of warm-acclimated mosquitoes could be fitted with a slope of -4, which was then used also for tolerance curves of warm-evolved and mosquitoes maintained at standard conditions to compare their vigour, sensitivity and severity using a likelihood ratio test (Table 1). All tested parameters showed significant differences between at least two groups. Warm-acclimated mosquitoes had the lowest vigour (95% CI: 12.38−13.39) and highest severity (95% CI: 9.44−11.29). Vigour of mosquitoes maintained at standard condition (95% CI: 16.23−17.43) and warm-evolved (95% CI: 15.85−17.06) mosquitoes was comparable. Warm-evolved mosquitoes also showed the lowest severity (95%CI: 12.95−14.35). Interestingly, both warm-acclimated (EC_50_ 95%CI 4.272 to not determined) and warm-evolved mosquitoes showed higher sensitivity than those kept at standard conditions (EC_50_ 95%CI 1.384−2.261) (Fig. 1D, Table S1). These results support the conclusion that exposing mosquitoes to hot increases their overall tolerance to a viral infection, but the length of the thermal challenge impacts vigour, severity and longevity. Multi-generational exposure to heat imposes no cost on CFAV infected mosquitoes.

**Table 1.**
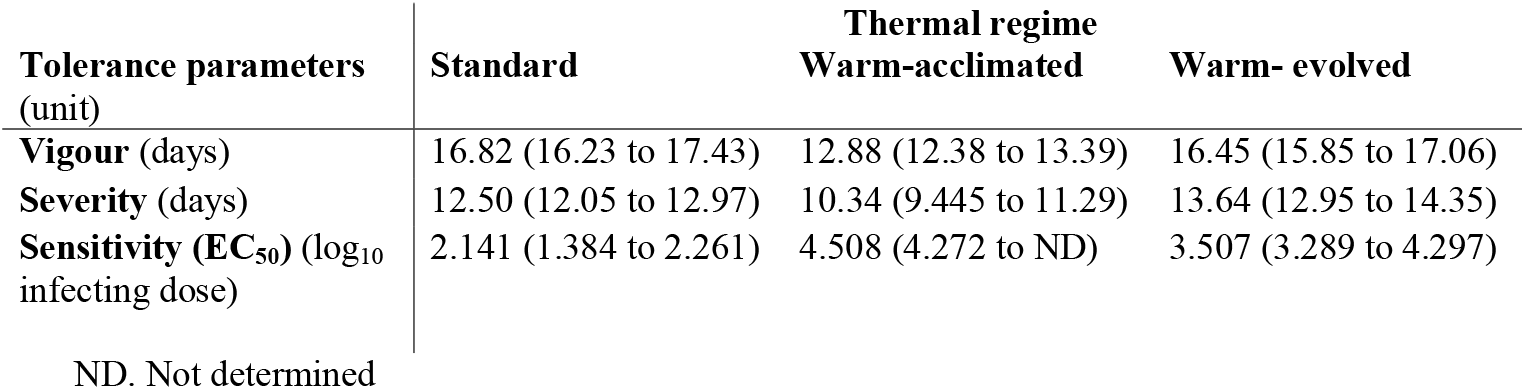
Tolerance curve parameters analysis. For each parameter the best-fit value and the 95% confidence interval (CI) are reported.

### Warm-acclimated *Ae. albopictus* have higher resistance

Next, we studied if exposure to heat alters *Ae. albopictus* resistance to CFAV. We decomposed resistance into two different phenotypes: prevalence of infection and viral load ^37^. We tested these traits 3- and 5-days post infection (dpi) in mosquitoes injected with the five viral doses used to assess tolerance. Prevalence was dose dependent. The two lower infecting doses, 10 and 10^2^ viral particles, resulted in a significantly lower prevalence compared to the higher three (Table S2). We also observed an effect of the length of the thermal challenge on the prevalence of infection. Warm-acclimated mosquitoes had lower prevalence than both warm-evolved mosquitoes and mosquitoes kept at standard condition regardless of the injected dose and dpi (warm-acclimated vs. mosquitoes kept at standard z.ratio=4.617, df=Inf, p<0.0001; warm-acclimated vs. warm-evolved z.ratio=4.939, df=Inf, p<0.0001). Additionally, warm-evolved mosquitoes showed lower prevalence of infection compared to mosquitoes kept at standard conditions at both 3 dpi (z.ratio=2.367, df=Inf, p=0.021) however this difference was lost at 5 dpi (z.ratio=1.425, df=Inf, p=0.1542). The length of the thermal challenge also influenced viral loads, in a time-dependent manner (Table 2). Warm-acclimated mosquitoes constrained viral infection more efficiently than both warm-evolved mosquitoes and mosquitoes kept at standard conditions within 3 dpi, independently of the injected concentration (Table S2, z.ratio=2.896, df=Inf, p=0.0004). This was a transient effect since at 5 dpi, mosquitoes from all regimens showed comparable viral loads (Table S2).

**Table 2.**
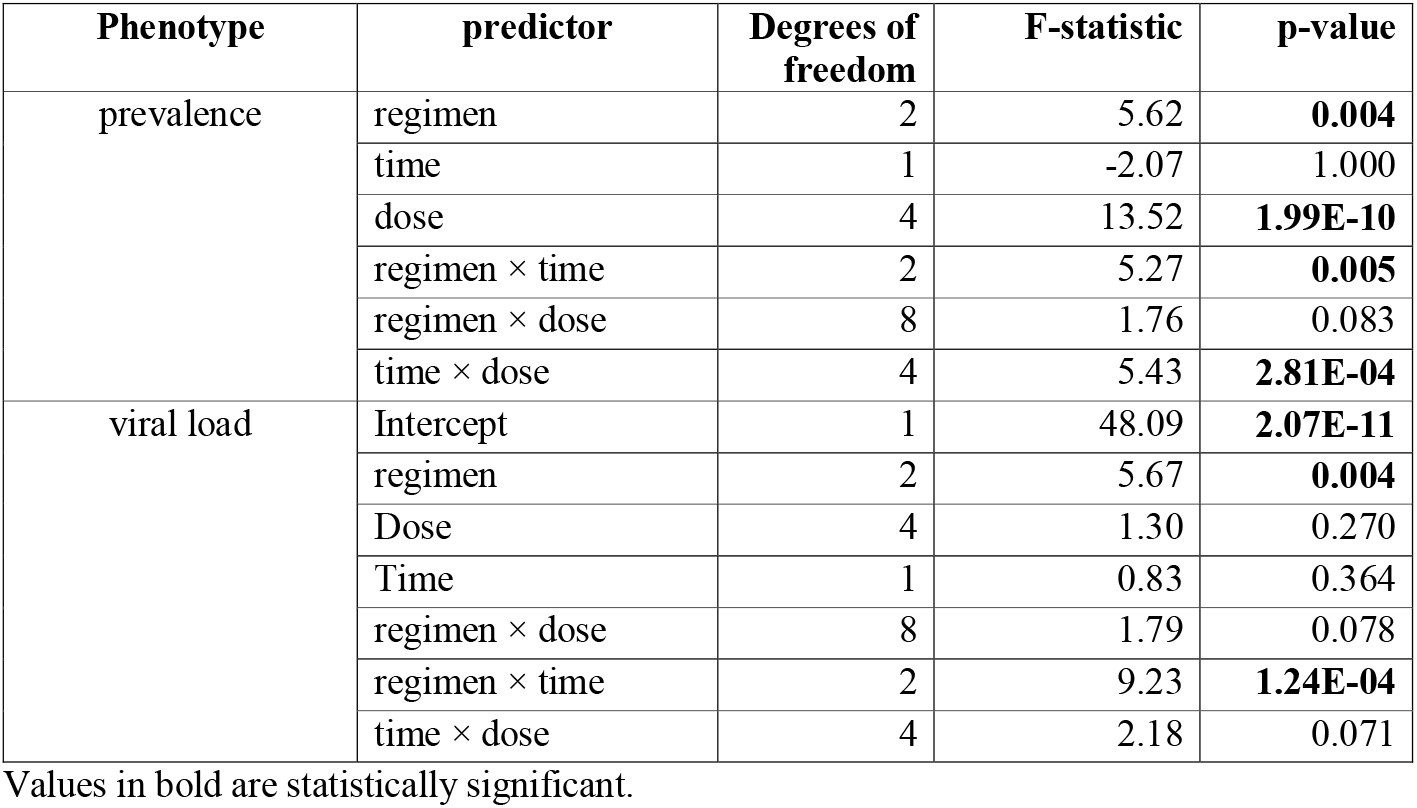
Effect of thermal regimens, CFAV infecting dose and infection period on *Ae. albopictus* resistance as assessed by analysis of variance (ANOVA) of linear models.

| Phenotype | predictor | Degrees of freedom | F-statistic | p-value |
| --- | --- | --- | --- | --- |
| prevalence | regimen | 2 | 5.62 | **0.004** |
|  | time | 1 | -2.07 | 1.000 |
|  | dose | 4 | 13.52 | **1.99E-10** |
| | regimen × time | 2 | 5.27 | **0.005** |
| | regimen × dose | 8 | 1.76 | 0.083 |
| | time × dose | 4 | 5.43 | **2.81E-04** |
| viral load | Intercept | 1 | 48.09 | **2.07E-11** |
|  | regimen | 2 | 5.67 | **0.004** |
|  | Dose | 4 | 1.30 | 0.270 |
|  | Time | 1 | 0.83 | 0.364 |
| | regimen × dose | 8 | 1.79 | 0.078 |
| | regimen × time | 2 | 9.23 | **1.24E-04** |
| | time × dose | 4 | 2.18 | 0.071 |

Values in bold are statistically significant.

### Warm-acclimated mosquitoes suffer extensive fitness costs from CFAV infection

Lastly, we examined the effect of heat exposure on *Ae. albopictus* and its influence on the fitness cost associated with CFAV infection. As a readout for the cost of infection, we used mosquito longevity and reproductive output, measured from five different parameters: percentage of sterile female, fecundity (*i*.*e*. number of eggs produced by each female), fertility (*i*.*e*. number of hatching eggs), percentage of non-viable eggs and progeny number per female (*i*.*e*. number of larvae/female). Linear and binomial generalized linear models were used to determine the effects of thermal regimens, CFAV infection, and their interaction on reproductive traits (Table S3). To decouple the effects of temperature and CFAV infection, we first compared the fitness of warm-acclimated, warm-evolved and mosquitoes kept at standard conditions without infection (Gray comparison in Fig. 4). We observed a significant reduction in female longevity (Fig. 4A), but higher fecundity, fertility and progeny/female in warm-acclimated vs. both warm-evolved and mosquitoes kept at standard conditions (Fig. 4 B-D, Table S3). We did not see significance differences in any of the tested parameters between warm-evolved and mosquitoes kept at standard conditions without infection (Fig. 4 A-D, Table S3). We saw that CFAV infection reduces longevity of warm-acclimated and mosquitoes kept at standard conditions, but not that of warm-evolved mosquitoes (Fig. 4A). CFAV infection also reduces fecundity of warm-acclimated mosquitoes (Fig. 4B, Table S3). Only the warm-acclimated mosquitoes showed an effect of injection on fertility and progeny (Fig 4C, D, Table S3). Standard mosquitoes also exhibited reduced fecundity, but this was due to the combined effect of the injection and CFAV infection. (Fig. 4B, Table S3). Warm-evolved mosquitoes showed no fitness effects due to either injection or CFAV infection (Fig. 4C-D, Table S3).

**Figure 4.**
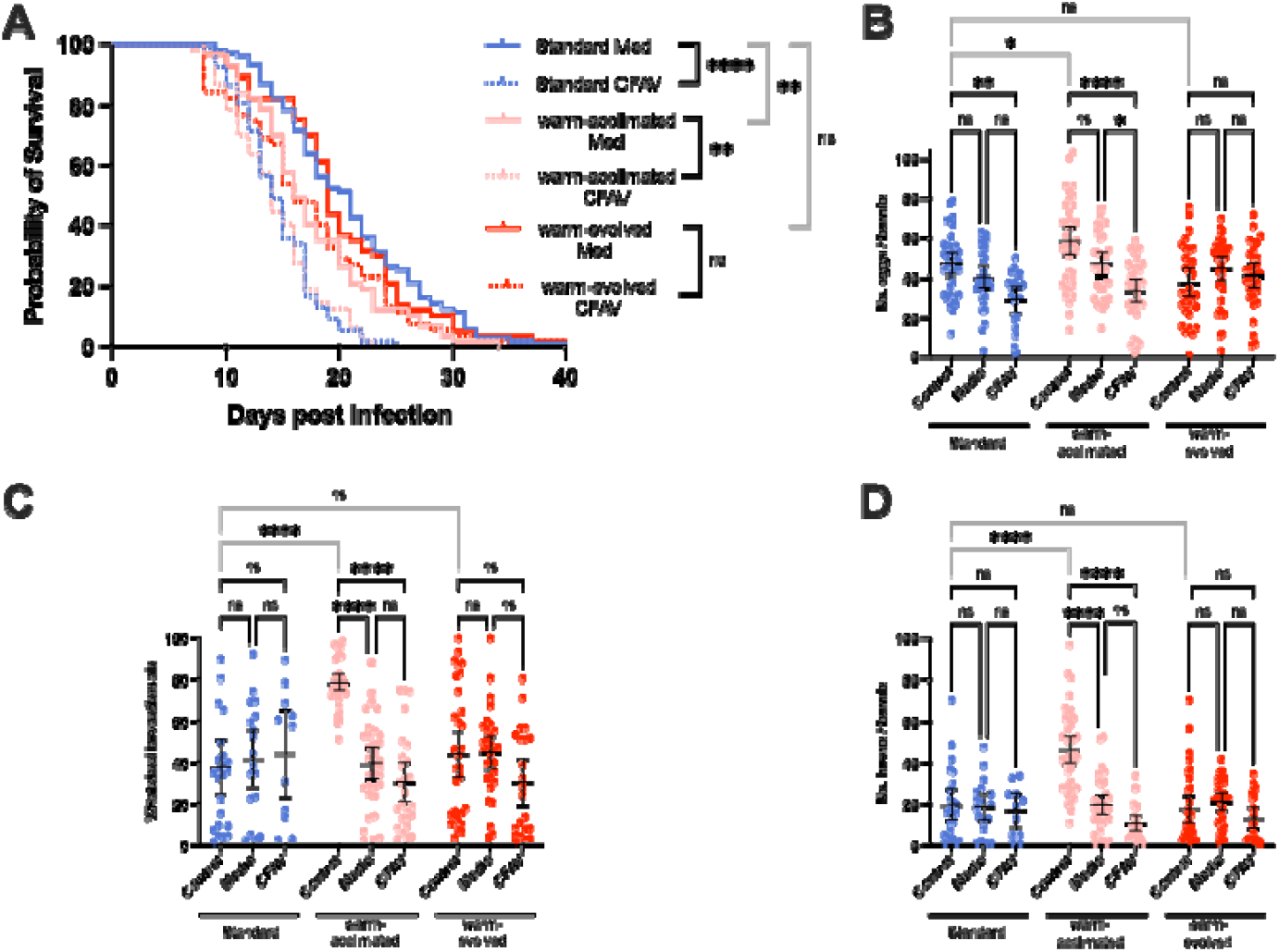
CFAV infection incurs fitness costs in warm acclimated mosquitoes. (**A**) longevity, (**B**) fecundity, (**C**) fertility (**D**) and number of larvae per female in warm-acclimated (pink), warm-evolved (red) and standard (blue) mosquitoes. Each point represents data from an individual mosquito; black solid lines are the mean and bars show the 95% confidence intervals (CI). In all the plots, control, medium and CFAV refer to data from non-injected, medium-injected, and mosquitoes infected with 10^5^ viral particles, respectively. Grey lines are used for between regimen and black lines for within regimen comparisons. Ns represents not significant, * P-value<0.05, ** P-value<0.01, *** P-value<0.001, and **** P-value<0.0001.

## Discussion

Transmission heterogeneity is an inherent property of arboviral diseases^2^. Biotic and abiotic factors, including natural variation of vector competence across mosquito populations and current human-driven climate change effects on both vectors and arboviruses are known to contribute to differences in transmission dynamics ^19,38-40^. The need of including these variables to frame more realistic models of transmission, especially at a local scale, is being increasingly appreciated ^41,42^. Our results highlight important shifts between resistance and tolerance to viral infection in *Ae. albopictus* dependent on the length of a thermal challenge, which have critical implications for understanding natural heterogeneity of mosquito vector competence under current climatic changes. One generational exposure to heat, which mimics a heatwave, results in fitness losses and increased resistance to viral infection. In contrast, mosquitoes conditioned to heat through several generations, which mimics current global warming, are as healthy as, if not healthier than, those not exposed to a thermal challenge and have increased tolerance to CFAV. Besides having important implications for understanding natural heterogeneity of mosquito vector competence, this immunological shift has also long-term impacts on both vectors and viruses because resistance and tolerance have drastically different ecological and evolutionary consequences ^16-18^. By directly fighting infection, resistance implies the emergence of an arm-race between the virus infective strategies and mosquito immunity that could foster molecular evolution through continuous selection of adaptation and counter adaptation ^43^. On the contrary, tolerance is not directed towards the pathogen, but encompasses strategies used to limit the damage caused by the infection ^44^. As such, tolerance should reduce selective pressure on both viruses and mosquitoes, resulting in different evolutionary trajectories. Ultimately, mosquito evolutionary trajectories will be influenced by the extent of the physiological and genetic factors shared between resistance and tolerance mechanisms because even if resistance and tolerance have opposite outcomes, they may be interlinked^45^. For instance, some resistance effectors, such as antimicrobial peptides or reactive oxygen species (ROS), may have direct or indirect toxic effects on the mosquitoes besides the pathogen, thus affecting its tolerance ^6^. Additionally, resistance can be energetically demanding ^44,46-49^ requiring appropriate energy management in relation to tolerance mechanisms, which manage host homeostasis during an infection.

While the concept of tolerance was introduced in the early ’30 ^50^, plant pathologists developed in the ’90 a biological and mathematical framework to distinguish effects of resistance and tolerance based on reaction norms that measure host longevity across a range of pathogen doses and the definition of resistance as “inverse of the pathogen burden” and tolerance as “the rate of change in host fitness at increasing viral loads” (i.e. the slope of the reaction norm) ^46,47,51^. These concepts are slowly breaching into vector biology based on the observation that arboviral infections are persistent in mosquitoes, with no apparent or limited fitness consequences ^52-54^. Moreover, recent studies in *Drosophila melanogaster* and *Ae. aegypti* showed the advantages of addressing insect response to pathogen infections accounting for tolerance ^11,36,55-57^. For instance, while in infection experiments using 1000 median tissue culture infectious dose (TCID_50_) of Drosophila C virus (DVC), *Dr. melanogaster* longevity appeared to depend on the presence of the *G9a* gene ^57^, when the role of *G9a* was studied using a dose-response curve approach, *G9a* knockout was shown to reduce fly longevity with a lower amount of virus with respect to control flies, indicating that this gene controls sensitivity to infection ^36^. Similarly, using a dose-response curve, blood meal was shown to have an immediate, but transient effect on resistance, and a more prolonged impact on tolerance to both bacterial and arboviral infections in *Ae. aegypti* ^55,56^. By applying a dose-dependent curve to study viral infection in *Ae. albopictus*, we identified previously unappreciated fitness costs of CFAV infection ^25^. Our results also showed that temperature influences CFAV prevalence, with one generational exposure to heat reducing viral prevalence. These results are important for potential applications of CFAV as a biological control agent to control arboviruses directly in their vectors ^58,59^. By highlighting the interplay between abiotic (*i*.*e*. temperature) and biotic (*i*.*e*. mosquito immunity) factors shaping CFAV prevalence in mosquitoes, our results could contribute in explaining the heterogenicity of its detection across mosquito field populations and caution on its use to elucidate mosquito population structure and dynamics ^28,29,59^.

Temperature is known to influence insect ability to mount an effective immune response against pathogens ^60^. Orthoflavivirus infection prevalence increases in *Ae. albopictus* and *Ae. aegypti* at high temperatures, alongside with dissemination and transmission rates ^5,61,62^. Exposure of adult mosquitoes to warmer temperatures, following development at temperate conditions, is also known to favour upregulation of heat shock proteins, which have pan-antiviral activity in mosquitoes ^63,64^. However, heat-shock proteins can be toxic and result in cellular damages at high concentrations, suggesting their upregulation is limited during an acute thermal stress ^64^. Owing to extensive cross-talks in the signalling cascades that regulates stress-related transcription factors ^65^, modification of the expression profile of antimicrobial peptides and other immune-related genes was also observed in mosquitoes exposed to a heat challenge ^4,66,67^. These results agree with our observation that one generational exposure to heat results in mosquitoes fighting viral infection, with lower tolerance with respect to warm-evolved mosquitoes. Because warm-acclimated mosquitoes had altered fitness compared to warm-evolved ones in the absence of infection, our results indicate that acclimation is a stressful condition, which conditions mosquito immunity. In contrast, warm-evolved mosquitoes did not suffer fitness costs with respect to mosquitoes reared at constant conditions either in presence or absence of CFAV infection, but showed less resistance in comparison to warm-acclimated mosquitoes. These results support the conclusions that short vs long term thermal challenges evoke different physiological mechanisms in mosquitoes, which require further investigations. These results also align with recent data from several mosquito species, including *Ae. albopictus* ^3^, which support the existence of thermal adaptation in mosquitoes ^68,69^. Thermal adaptation includes profound and lasting behavioural and physiological changes, which allow a species to develop, survive and reproduce under otherwise unfavourable conditions such as those associated with long-term global warming ^32^.

To conclude, we highlight important shifts in tolerance and resistance to viral infection dependent on the length of the thermal challenge, which underlines the complexity of climate change effects on mosquito biology and arboviral transmission dynamics.

## Materials and methods

### 1. Mosquito rearing

We used the Foshan laboratory population ^70,71^, which we maintain in a Binder KBWF climatic chamber under a constant temperature of 28°C, 70±5% relative humidity and a 12:12 hours light/dark photoperiod. We rear larvae at the density of approximately 200 individuals in one litre of distilled water in BugDorm plastic pans (19×19×6cm). We feed larvae daily with 10 pellets of Tetra Goldfish Gold Colour fish food (Tetra Werke, Germany) until pupation. We keep adult mosquitoes in 30 cm^3^ BugDorm cages and give them *ad libitum* access to 20% sucrose solution. We offer defibrinated mutton blood (Biolife Italiana) to females 7−10 days post emergence using a Hemotek feeding apparatus.

### 2. Thermal challenge

We exposed Foshan mosquitoes to a thermal challenge consisting of 14 h light at 32°C and 10 h dark at 26°C. We chose 32°C because it corresponds to the average T_a_ registered in the summer of 2021 in northern Italy and was previously shown to be a thermal condition above the optimum for Ae. albopictus ^3^. We designed three experimental groups exposed to two different thermal conditions: 1. Standard: mosquitoes reared under laboratory standard conditions; 2. warm-acclimated: mosquitoes reared under thermal challenge for one generation; 3. warm-evolved: mosquitoes reared under thermal challenge for ten generations. We established three replicates for each experimental group starting each with 1000 eggs. This number of eggs was maintained across generations for the warm-evolved mosquitotes.

### 3. Viral preparation

We obtained CFAV (Rio Piedras 2002, Ref-SKU: 001v-EVA68) from Ronald Van Rij (Radboud University, Nijmegen, NL). We propagated the virus in C6/36 cells maintained at 28°C on Leibovitz’s L-15 Medium (Gibco, REF:11415-049) with 2% FBS (Gibco, REF: 16000-044 Lot: 2421078RP). Three T75 tissue cultured flasks at an 80% confluency were used to start viral propagation. We used one of the three flasks to determine the number of cells present at the time of infection to calculate the multiplicity of infection (MOI). A second flask was infected at MOI of 0.1 (*i*.*e*. 1 viral particle per 10 cells). After five days, we clarified the media by centrifugation at 4000 × *g* for ten minutes at 4°C. Viral stock was concentrated with an AMICON Ultra-15 100k centrifugal filter (Millipore, UFC910008) using a swing bucket rotor and centrifugation at 4000 × *g* for 30 minutes. We quantified the viral stock by a focus forming assay and stored it at -80 °C. A third flask was kept at the same conditions of the infected one, including collection and storage of its media, which was used to inject mock infected mosquitoes.

### 4. Viral infection

Three to eight days old females were used in infection experiments. All infections were performed between 2 to 5 p.m. to reduce biases due to the circadian rhythm ^72^. We injected mosquitoes intrathoracically with 50 nL of CFAV or medium from C6/36 cells using the Nanoject III (Durmond). The volume of the injected virus was kept constant through viral concentrations, which were generated in RPMI-1640 media (2% FBS) when needed. Prior the injection, we cold anesthetized mosquitoes and separated them in three groups. First group is the control (Ctr); these females were not injected, but were kept on ice for the same time as injected mosquitoes. The second group was mock infection (Med), including females injected with media from C6/36 cells. The third group was. CFAV infected (CFAV) females, meaning females injected with CFAV. After infection, we returned all mosquitoes to the same rearing condition where they were raised until needed for further experiments.

### 5. Tolerance

We quantified tolerance by measuring longevity of mosquitoes infected with five increasing viral concentrations, such as 10, 10^2^, 10^3^, 10^4^, 10^5^ viral particles ^34^. For each concentration we used 20 females, along with a group of 20 mosquitoes mock infected with media and a group of 20 mosquitoes that were not injected. After infection, we kept mosquitoes in 266 mL cardboard cups, added a cotton ball with water and a cotton ball with 20% sugar. Mosquitoes that died within 3 days post infection were not included in survival analyses because we assumed their death was caused by injection-related injuries. This experiment was repeated three times for mosquitoes kept at standard condition or exposed to thermal challenge for one or ten generations.

### 6. Resistance

We infected mosquitoes with the same viral concentrations used to determine tolerance and we measured viral genomes in mosquitoes three- and five-dpi, through quantitative polymerase chain reaction (qPCR).

We extracted RNA using 500 µL of TRIzol Reagent (Invitrogen, Ref: 15596018) per mosquito, followed the manufactures’ manual and resuspended the final pellet in 20 µL of Ultra-pure water. Then, 10 µL of the total RNA was used to produce cDNA with the GoScript Reverse Transcription Mix (Promega) following manufacturer’s recommendation. We used the QuantiNova SYBR Green PCR kit (Qiagen) for qPCR and primers targeting the *NS3* viral gene ^24^. The 2-_ΔΔ_Ct method was used to calculate viral loads relative to the housekeeping *Rpl34 Ae. albopictus* gene ^73^. We infected 12 mosquitoes per concentration and collected 6 mosquitoes per timepoint; we repeated the experiment three times for mosquitoes kept at standard condition or exposed to thermal challenge for one or ten generations.

### 7. Mosquito fitness

We infected three to eight old females with 10^5^ viral particles. After infection, females were transferred to separate cages in groups of 50 and 10 three-to-eight-day old males were added to each cage for mating. Seven days post injection, we offered a blood meal to females. We transferred engorged females to cardboard cups in groups of 12-20 for two days and then we transferred them to individual plastic cups. We added filter paper and water at the bottom of the cup as oviposition site, mesh on top and cotton balls to provide sucrose solution. Seven days after blood feeding, we counted the number of eggs laid per female, as a measure of their fecundity. We dried eggs for two days before hatching them. The percentage of hatched eggs over the total number of eggs laid per female was measured as fertility. We also counted the number of larvae resulting from eggs laid by each female as progeny production. We further recorded the percentage of females that did not oviposit egg and the number of females that produce non-fertile eggs as sterility and infertility, respectively.

### 8. Statistical analysis

Analyses of resistance, survival and reproductive output were done with R version 4.3.3 (The R Foundation for Statistical Computing, 2024). They all included the temperature treatment (standard, warm-acclimated and warm-evolved), injected dose (10 to 10^5^ CFAV viral particles) and their interaction as fixed factors. Analysis of resistance included days post injection (3 dpi and 5 dpi) and its interaction with all other variable as an additional fixed factor. Significance of each factor was calculated with the function ANOVA from the car package version 3.1-2^74^. To estimate the correlation coefficients between viral dose and treatment, each model was reduced to a minimal model by removing non-significant interactions. Whenever a significant interaction remained, we used a type III ANOVA; otherwise, we used a type II. We used the functions emmeans and pairs from the emmeans package version 1.10.0 ^75^ for post-hoc analysis, adjusting p-values with the Tukey method. To assess tolerance, mosquito survival post injection was analyzed using a Cox proportional hazards model (function coxph from the survival package version 3.5-8). Model assumptions were tested using the cox.zph function (survival package version 3.5-8) ^76^.

We used GraphPad Prism (Version 10.2.0) to fit the four parameter logistic model as described by the following equation: Y=Bottom + (Top-Bottom)/(1+10^((LogEC_50_-X)×HillSlope)), where EC_50_ is the concentration of CFAV that reduces mosquito lifespan by half (between Bottom and Top). HillSlope is the steepness of the dose-response curve, and Top and Bottom are plateaus in the units of the Y axis.

Prevalence of viral infection was analyzed with a Bayesian generalized linear model (function bayesglm from the arm package version 1.13-1)^77^ with binomial distribution of errors. This approach was chosen to account for the observed quasi-separation of data for prevalence. Non-zero viral load were analyzed with a linear mixed-effect model with a normal distribution of errors (function Lmer from the lme4 package version 1.1-35.1)^78^. To reach normality of the residuals, we performed a boxcox transformation on the log10(viral titre) values using the function boxcox from the MASS package version 7.3-60.0.1^79^.

The proportion of female that laid did not laid eggs (sterile female) and the ensuing proportion of laid eggs that did not hatched (non-viable eggs) were analyzed respectively with a generalized linear mixed-effect model (function Glmer from the lme4 package version 1.1-35.1)^78^ and Bayesian generalized linear model (function bayesglm from the arm package version 1.13-1)^77^ with a binomial distribution of errors. Number of eggs laid, percentage of eggs hatched and number of progeny per female were analyzed with linear mixed-effect models with a normal distribution of errors (function Lmer from the lme4 package version 1.1-35.1)^78^. Whenever relevant, model assumptions were assessed using the Dharma package (version 0.4.6)^80^. All graphs were generated using GraphPad Prism (Version 10.2.0).

